# ERF1 inhibits lateral root emergence by promoting local auxin accumulation with altered distribution and repressing *ARF7* expression

**DOI:** 10.1101/2023.03.02.530895

**Authors:** Pingxia Zhao, Jing Zhang, Siyan Chen, Zisheng Zhang, Guangyu Wan, Jieli Mao, Zhen Wang, Shutang Tan, Chengbin Xiang

## Abstract

Lateral roots (LRs) are crucial for plants to sense environmental signals in addition to water and nutrient absorption. Auxin is key for LR formation, but the underlying mechanisms are not fully understood. Here we report that *Arabidopsis* ERF1 inhibits LR emergence by promoting local auxin accumulation with altered distribution and regulating auxin signaling. Loss of *ERF1* increases LR density compared with the wild type, whereas *ERF1* overexpression causes the opposite phenotype. ERF1 enhances auxin transport by upregulating *PIN1* and *AUX1*, resulting in excessive auxin accumulation in the endodermal, cortical, and epidermal cells surrounding LR primordia. Furthermore, ERF1 represses *ARF7* transcription, consequently affecting the expression of cell wall remodeling genes that facilitate LR emergence. Together, our study reveals that ERF1 integrates environmental signals to promote local auxin accumulation with altered distribution and repress *ARF7*, consequently inhibiting LR emergence in adaptation to fluctuating environments.

**Highlights:**

- ERF1 functions as a negative regulator of lateral root emergence
- ERF1 enhances rootward and shootward auxin transport by directly upregulating the expression of *PIN1* and *AUX1*, resulting in high local auxin accumulation and abnormal auxin distribution in the endodermal, cortical, and epidermal cells overlying lateral root primordia
- ERF1 represses the transcription of *ARF7* and cell wall remodeling genes in lateral root emergence

## Introduction

Lateral roots (LRs) are crucial for plants to forage nutrients from soil and sense underground signals.^1^ LRs originate primarily from the pericycle cells, where a series of asymmetric cell divisions form a dome-shaped primordium.^2–4^ LR primordium (LRP) must traverse the endodermal, cortical, and epidermal cell layers to develop into LR. The whole process of LR formation is artificially divided into 8 stages.^5, 6^

Auxin is a master regulator required for LR development, to which endogenous and environmental signals converge.^7–9^ To date, numerous studies have shown that LR developmental events from priming to initiation, patterning, and emergence are regulated by auxin. LR initiation is a phase encompassing specified lateral root founder cells (LRFCs) undergoing nuclear migration and several rounds of anticlinal asymmetric divisions at regular intervals along the primary root.^7^

Auxin accumulation in root pericycle cells is one of the earliest events for founder cell (FC) specification to give rise to LR initiation.^10, 11^ The auxin maximum is observed in the protoxylem cells of the basal meristem and prior to the actual initiation event.^12^ How the auxin maximum is built up in protoxylem cells is not completely understood. The local auxin maximum is generally considered to be created and maintained through the activation of auxin biosynthesis and transport. The establishment of a local auxin gradient through auxin transport is required for organ formation.^13, 14^ Recent studies have demonstrated that auxin transport from lateral root cap (LRC) cells to the inner tissue of the elongation zone causes an auxin maximum response in the oscillator zone (OZ). Subsequently, the auxin maximum is attenuated before increasing gradually again in the OZ, which contributes to the specification of xylem pole pericycle (XPP) cells for LR initiation and spacing.^15–18^ The membrane- localized auxin efflux carrier PIN (PIN-FORMED)-mediated local auxin gradient is essential for LRP formation.^13^ Multiple *pin* mutants (*pin1 pin3*, *pin1 pin4 pin7* or *pin1 pin3 pin4 pin7*) show defects in LRP development.^13, 19, 20^ Similarly, the members of the ABC-B/multidrug resistance/P-glycoprotein (ABCB/MDR/PGP) subfamily also act as auxin efflux carriers. ABCB1 and ABCB4 facilitate auxin entry into the shootward stream from the root apex, but their auxin-exporting capability is lower than that of PINs.^21–24^ Moreover, the auxin influx carrier AUX1, which mediates shootward auxin transport throughout the LRC and epidermis, is important for LR positioning, initiation, and emergence.^15, 16, 25, 26^ The *aux1* mutant displays reduced auxin levels in LRC cells, which inhibits prebranch sites and LR formation.^3, 16^ *LIKE AUX1* (*LAX*) *3* is induced in epidermal and cortical tissue by auxin and is involved in auxin distribution in cells overlaying LRP, promoting LR emergence.^27, 28^ Additionally, auxin passive diffusion via plasmodesmata has been demonstrated to contribute to auxin distribution.^29, 30^ Recently, water and auxin fluxes in the root basal meristem through plasmodesmata were reported, establishing a symplastic pathway to deliver auxin from the epidermis to the pericycle.^31^

The effect of auxin on plant growth is determined by auxin levels and distribution; therefore, a particular pattern of auxin distribution is required for root development. The distribution of endogenous auxin in LRFCs is altered by the inhibition of basipetal auxin transport, which blocks LR initiation.^7, 32^ Enhanced auxin biosynthesis or signaling in specific cell types of the epidermis, cortex, and endodermis inhibits root growth.^33^ The mutant *solitary root* (*slr-1*) exhibits strongly inhibited expression of *DR5pro:GUS* in the cortex and epidermis but not in the endodermis after auxin treatment,^27^ indicating that auxin distribution and response in the endodermis are important for LR emergence. Furthermore, *GNOM*/*EMBRYO DEFECTIVE30* (*EMB30*) mediates the establishment of the auxin response maximum in LRFCs, probably by regulating local and global auxin distribution.^34^ These studies show that auxin maximum establishment and its precise positioning are critical for LR initiation.

Auxin signaling is also involved in the regulation of the timing, pattern, and location of cell differentiation during LR formation. Auxin signaling modules composed of Aux/IAA transcriptional repressors and auxin response factors (ARFs) mediate auxin signaling to regulate different stages of LR formation. Among the ARFs, ARF7 and ARF19, which act in the IAA14/28-dependent auxin signaling module, coregulate the expression of the *LBD16*, *LBD18*, *LBD29*, and *LBD33* genes involved in LRFC polarization and LR initiation.^35, 36^ In the gain-of-function mutant *SLR/IAA14* (*slr-1*) and double mutants *arf7 arf19* and *lbd16 lbd18*, LR initiation is severely inhibited.^7, 35, 37^ LR emergence is the process by which LR grows through the endodermal, cortical and epidermal cell layers of the primary root, which is associated with local cell wall loosening. Therefore, cell wall remodeling (CWR) also contributes to LR emergence. Auxin-regulated changes in the cell wall properties of the primary root cells overlying LRP are required for LR formation.^38, 39^ Auxin induces *LAX3* expression in the cortex and epidermis overlying LRP, leading to the upregulation of the CWR genes *AIR3*, *XTR6*, and *PG*, which facilitates LR emergence.^27^ The SHY2- mediated auxin response in the endodermis triggers changes in cell volume and shape, which are essential for LRP initiation.^39^ *EXP17,* encoding a cell wall-loosening factor, is upregulated by LBD18 to promote LR emergence during the auxin response.^40^ Furthermore, auxin-dependent INFLORESCENCE DEFICIENT IN ABSCISSION (IDA)-HAESA (HAE)/HAESALIKE2 (HSL2) signaling controls LR emergence by altering the expression of CWR genes.^41^

Endogenous and exogenous signals are integrated into auxin to regulate LR formation. It is well known that auxin induces the formation of LRP, but how high auxin levels caused by auxin transport affect LR emergence is not well understood. We previously reported that *Arabidopsis* ERF1 integrates ethylene signaling into auxin biosynthesis by upregulating *ASA1* expression and thereby increasing auxin accumulation in the root tip, leading to the inhibition of primary root elongation.^42^

Here, we show that ERF1 functions as a negative regulator of LR emergence. We demonstrated that ERF1 enhanced auxin transport by directly upregulating the expression of *PIN1* and *AUX1*, thereby causing high local auxin accumulation in the endodermis, cortex and epidermis overlying LRP. Moreover, ERF1 transcriptionally repressed *ARF7* and consequently downregulated CWR genes, ultimately inhibiting LR emergence. Therefore, our results support that ERF1 plays an important role in integrating environmental signals into auxin signaling to regulate LR emergence.

## Results

### ERF1 inhibits LR emergence

A previous study showed that *Arabidopsis* ERF1 integrates ethylene signaling into auxin biosynthesis to regulate primary root elongation.^42^ To investigate the function of *ERF1* in LR development, we generated a knockout mutant, *erf1-2,* with an adenine (A) insertion in the codon of the 62^nd^ amino acid resulting in premature termination from AGT to TAG at the 65^th^ codon of *ERF1* mRNA translation by the CRISPRLCas9 system (Figure S1A). The RNAi knockdown line (*RNAi-1*) and overexpression lines as described ^42^ were confirmed by analyses of *ERF1* expression (Figure S1B), and then we performed LR development assays. The *RNAi-1* and *erf1-2* lines exhibited more LRs than the wild type (WT) when grown on MS medium, whereas the overexpression (OX) lines showed strong suppression of LRs (Figures 1A and 1B). Similar results were observed for LR density (Figure 1C).

**Figure 1.**
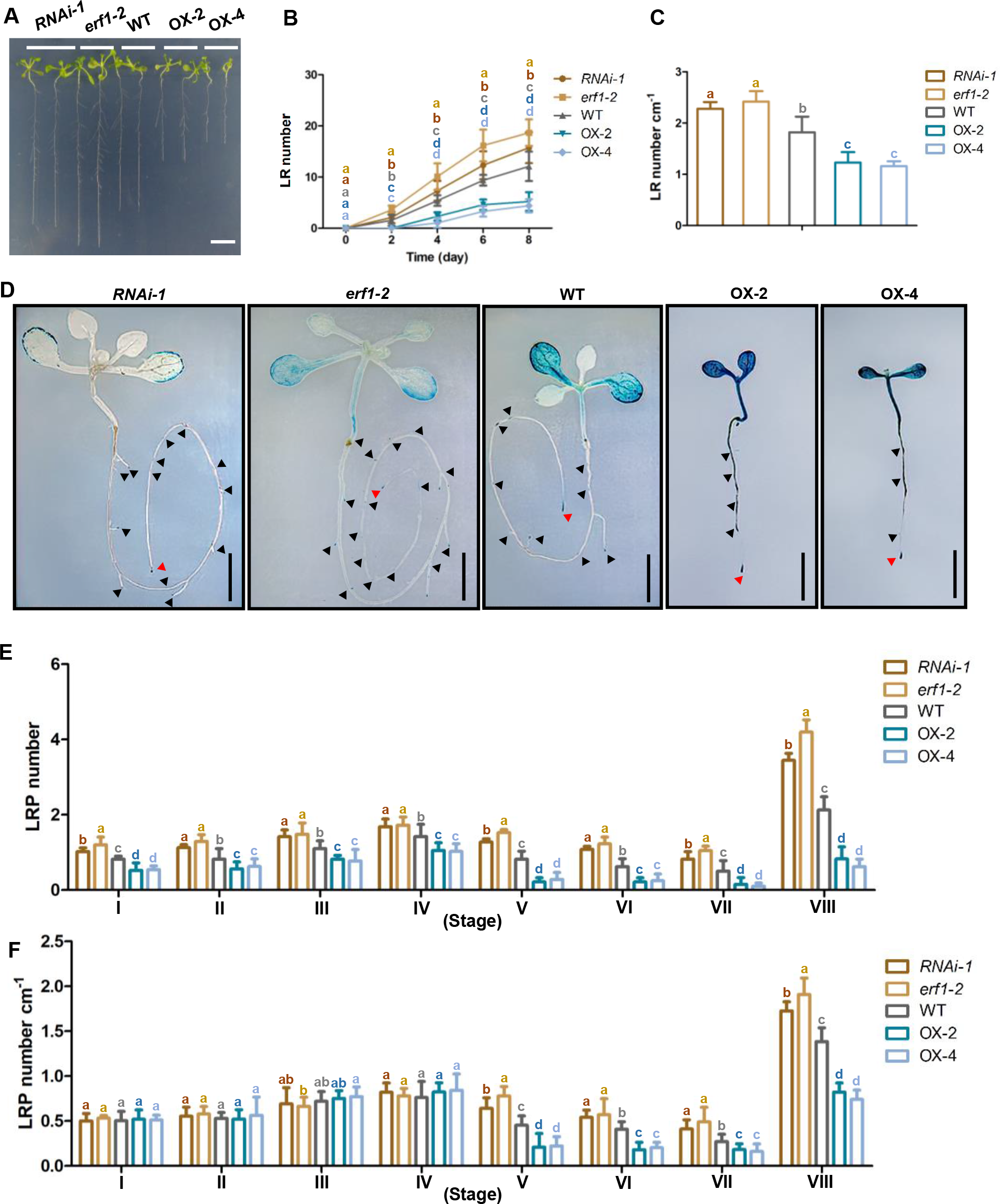
ERF1 inhibits LR emergence. (A-C) Lack of *ERF1* increases LR number. Seeds of *RNAi-1*, *erf1-2*, WT, OX-2, and OX-4 lines were germinated on MS medium for 5 days, then the seedlings were transferred to MS medium and vertically grown for 8 days before photographs were taken (A). The LR number was counted every other day for 8 days (B). The LR number cm^-1^ was calculated at day 8 (C). Values are mean ± SD (n=3 replicates, 50 seedlings/replicate). Different letters indicate significant differences by one-way ANOVA (P < 0.05). Bar=1 cm. (D) Phenotype of LR emergence in *erf1*, WT, and OX lines. Ten-day-old seedlings of *RNAi-1*, *erf1-2*, WT, OX-2, and OX-4 in the *DR5::GUS* background were stained for GUS and the locations of LRP were indicated by black triangles. Bar=1 cm. (E and F) The LRP number at given developmental stages. Seeds were germinated and vertically grown on MS medium for 10 days before the LRP number (E) was counted and LRP number cm^-1^ (F) was calculated. Values are mean ± SD (n=3 replicates, 50 seedlings/replicate). Different letters indicate significant differences by one-way ANOVA (P < 0.05).

To explore which stages of LR development were affected by *ERF1*, we generated *DR5::GUS* marker lines in the *erf1*, WT, and OX backgrounds by crossing and counted LRP in these lines. Consistent with the results shown in Figure 1B, the *RNAi-1* and *erf1-2* lines displayed more LRP than the wild type at stages I-VIII, especially at stage VIII, while the overexpression lines displayed a considerable decrease in LRP number compared with the wild type (Figures 1D and 1E). However, LRP density showed no significant difference among all the lines at stages I-IV, whereas it was higher in the *RNAi-1* line and *erf1-2* mutant but lower in the OX lines at stages V-VIII (Figure 1F). These results indicate that ERF1 negatively regulates LR emergence.

### ERF1 promotes auxin transport

ERF1 is known to increase auxin accumulation in primary roots by activating *ASA1*.^42^ To determine whether ERF1 modulates auxin levels in LR emergence, we checked auxin accumulation in the LRP of *erf1*, WT, and OX lines with the *DR5::GUS* background by GUS staining. The GUS signal was reduced in the LRP of *RNAi-1* and *erf1-2* mutants but markedly increased in the OX lines compared with that of the wild type. In OX roots, strong GUS signals were observed in LRP and the epidermal, cortical and endodermal cells overlying LRP at stages I-VIII, which were not visible in the wild type or *erf1* mutants (Figure 2A). The relative GUS intensity is shown in Figure 2B. These results show that ERF1 increases auxin accumulation and alters the auxin distribution pattern in the region where LRP occurs.

**Figure 2.**
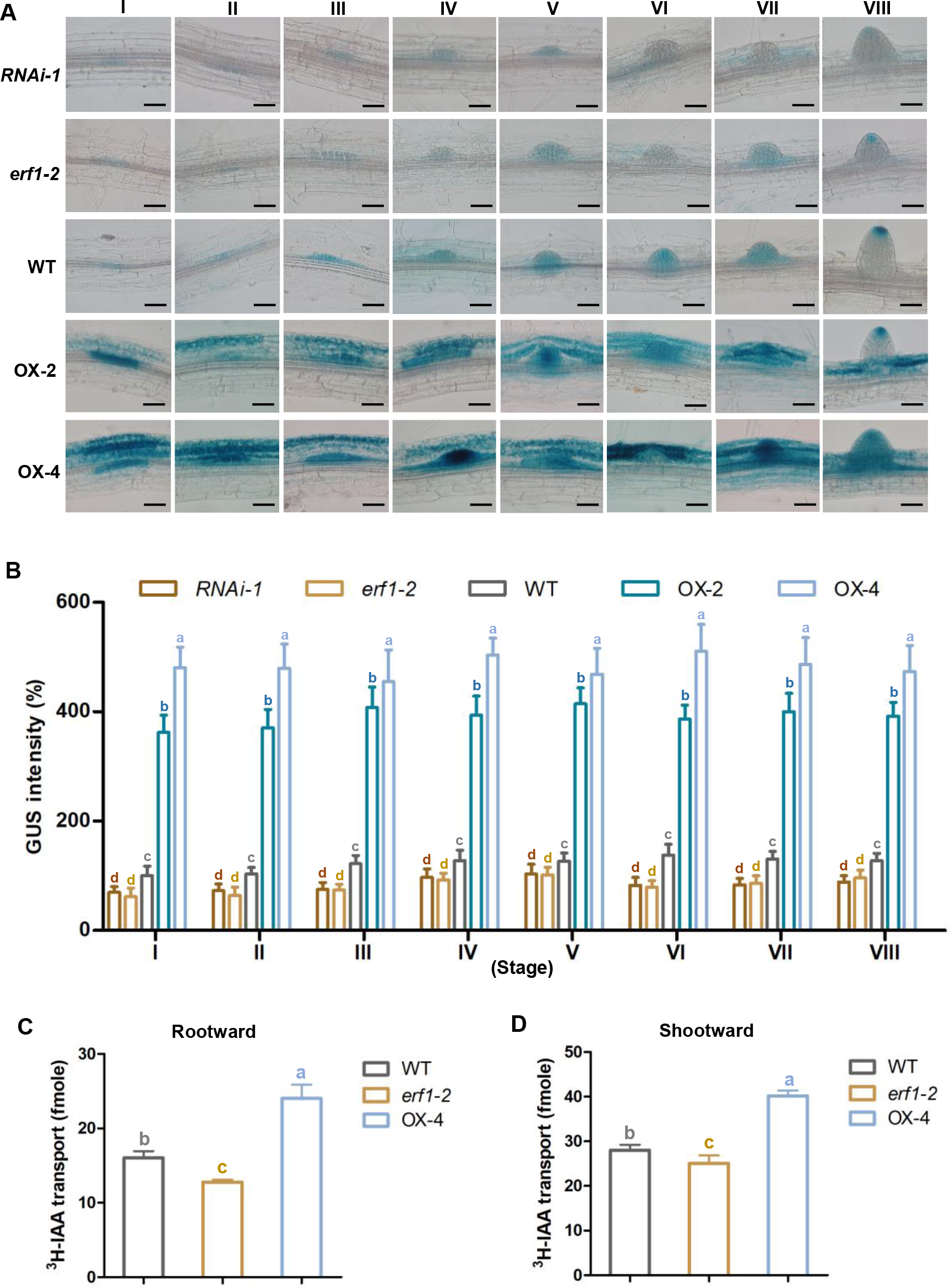
ERF1 enhances rootward and shootward auxin transport. (A and B) Auxin accumulation in LR development. Seeds of *RNAi-1*, *erf1-2*, WT, OX-2, and OX-4 lines in *DR5::GUS* background were vertically germinated on MS medium for 10 days, then seedlings were stained in GUS solution for 5 hours. The photographs of stages I to VIII were taken by microscope (A). Bar=50 μm. GUS signal intensity was quantified with ImageJ software (B). Values are the mean ± SD (20 images/genotype). Different letters indicate significant differences by one-way ANOVA (P < 0.05). (C and D) Auxin transport assays. Seven-day-old seedlings of *erf1-2*, WT, and OX-4 lines were used for auxin transport assays with ^3^H-IAA as described in the Methods. Rootward (C) and shootward (D) ^3^H-IAA transport were quantified. Values are mean ± SD (n=3 replicates, 30 seedlings/replicate). Different letters indicate significant differences by one-way ANOVA (P < 0.05).

To explore whether the enhanced auxin levels in OX lines were caused by auxin transport, we measured auxin transport in the seedlings of *erf1-2*, WT, and OX-4 by a ^3^H-IAA feeding experiment. Rootward and shootward auxin transport of the *erf1-2* line was decreased compared with that of WT but elevated in the OX-4 line (Figures 2C and 2D). These results suggest that ERF1 enhances rootward and shootward auxin transport, which might contribute to high local auxin accumulation and alter auxin distribution in the epidermis, cortex and endodermis overlying the LRP of the OX-4 line.

To further verify the involvement of auxin transport in LR emergence, we treated seedlings of *erf1*, WT, and OX lines with the IAA efflux inhibitor *N*-1- naphthylphthalamic acid (NPA).^43, 44^ LR emergence was severely inhibited in all genotypes when treated with NPA. However, it was more severely inhibited in the WT and OX lines than in the *erf1* lines (Figures 3A-3C). We also examined auxin accumulation in the LRP of *erf1-2*, WT, and OX-4 lines under NPA treatment. The GUS signals were distinctly suppressed in the LRP of the WT and OX lines but increased in their root tips. In contrast, GUS signals were almost unchanged in the LRP and root tip of the *erf1-2* mutant (Figures 3D and S2). Taken together, these data indicate that ERF1-modulated auxin accumulation is dependent on auxin transport.

**Figure 3.**
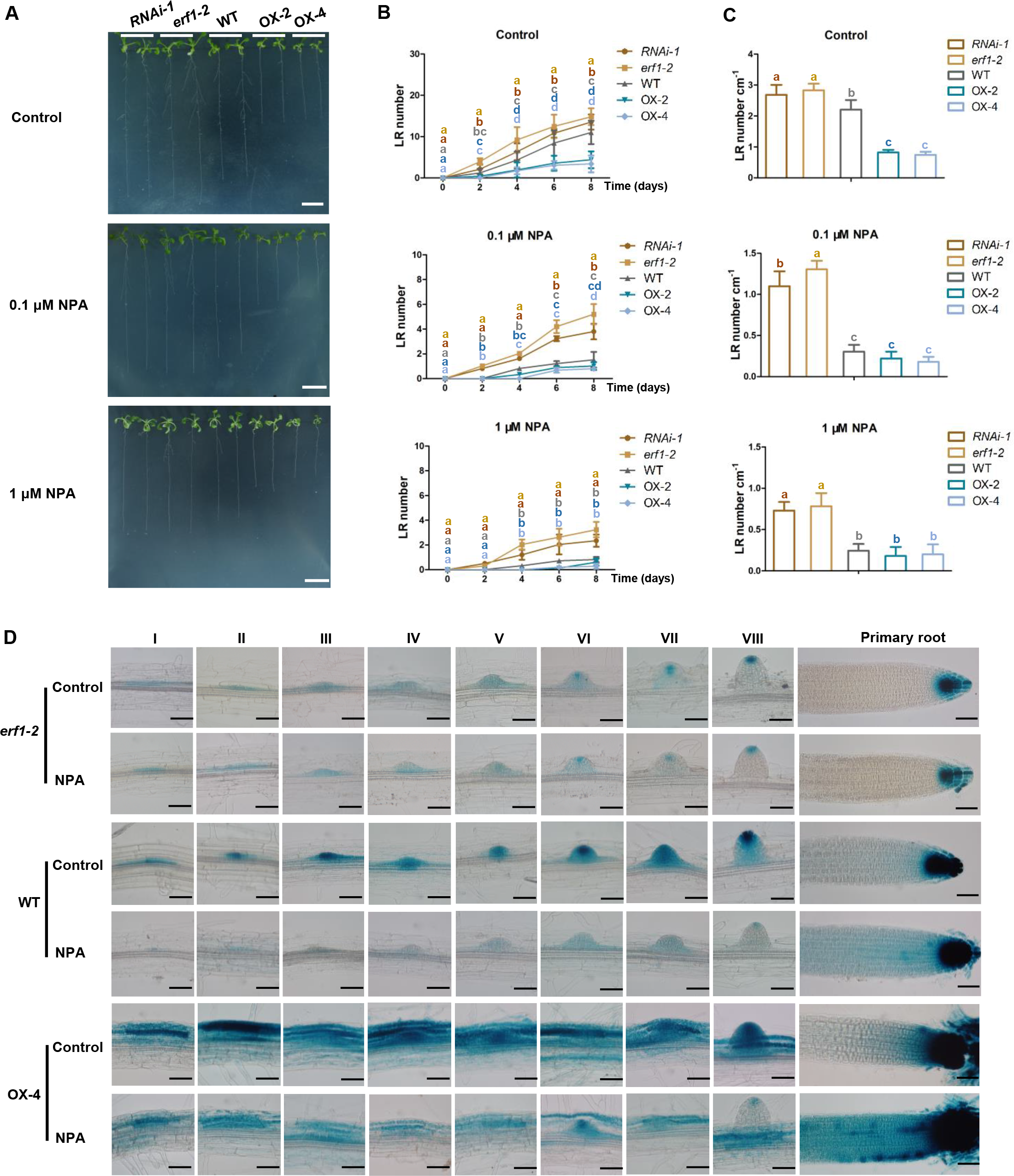
NPA treatment eliminates the difference in LR number and density between WT and OX lines. (A-C) NPA treatment severely reduces LR emergence in *erf1*, WT, OX lines. Seeds of *RNAi-1*, *erf1-2*, WT, OX-2, and OX-4 lines were germinated on MS medium for 5 days, and then seedlings were transferred to MS medium without (Control) or with 0.1 and 1 µM NPA and grown for 8 days before photographs were taken (A). LR number was counted every other day for 8 days (B). LR number cm^-1^ was calculated at day 8 (C). Values are mean ± SD (n=3 replicates, 50 seedlings/replicate). Different letters indicate significant differences by one-way ANOVA (P < 0.05). Bar=1 cm. (D) NPA treatment reduces auxin accumulation in LRP. Seeds of *erf1-2*, WT, and OX-4 lines in *DR5::GUS* background were germinated on MS medium for 5 days, then transferred to MS without (Control) or with 1 μM NPA and grown for 2 days. Seedlings were incubated in GUS staining solution for 8 hours before the photographs of LRP from stages I to VIII as well as the primary tips were taken by microscope. Bar=50 μm.

### ERF1 directly regulates the transcription of *PIN1* and *AUX1*

To address how *ERF1* regulates auxin transport, we measured the transcript levels of *PIN1*, *PIN2*, *PIN3*, *PIN4*, *PIN7* (auxin efflux transporters); *AUX1*, *LAX1*, *LAX2*, *LAX3* (auxin influx transporters); and *PID*, *WAG1*, *WAG2* (PIN phosphorylation kinase) in *erf1-2*, WT, and OX-4 lines by qRTLPCR. *PIN1*, *PIN3*, *PIN4*, *PIN7*, *AUX1*, *LAX2*, and *LAX3* were significantly downregulated in the *erf1-2* mutant compared with the WT after 7 and 10 days of growth, whereas *PIN1*, *AUX1*, *LAX1*, and *LAX3* were upregulated in the OX-4 line (Figure S3). Therefore, ERF1 is likely involved in regulating auxin transport by modulating the expression of the genes related to auxin transport.

To link ERF1 with these auxin transport-related genes, we analyzed the potential binding sites (GCC-box) of ERF1 in the promoter of auxin transport-related genes and found that *PIN1* and *AUX1* contained one GCC-box in each 2.5 kb promoter region (Figures 4A and 4B). ChIPLqPCR assays were carried out using WT and *35S::HA- ERF1* transgenic plants. The promoter fragments containing the GCC-box of *PIN1* and *AUX1* were significantly enriched in *35S::HA-ERF1* transgenic plants (Figures 4C and 4D). Electrophoretic mobility shift assays confirmed that ERF1 was able to directly bind to the promoters of *PIN1* and *AUX1* (Figures 4E and 4F). Additionally, we conducted transient expression assays in tobacco leaves and demonstrated that ERF1 was able to activate the expression of the *PIN1* and *AUX1* promoters fused with LUC (Figures 4G and 4H). Together, these results show that ERF1 can activate the expression of *PIN1* and *AUX1* by binding to their promoters.

**Figure 4.**
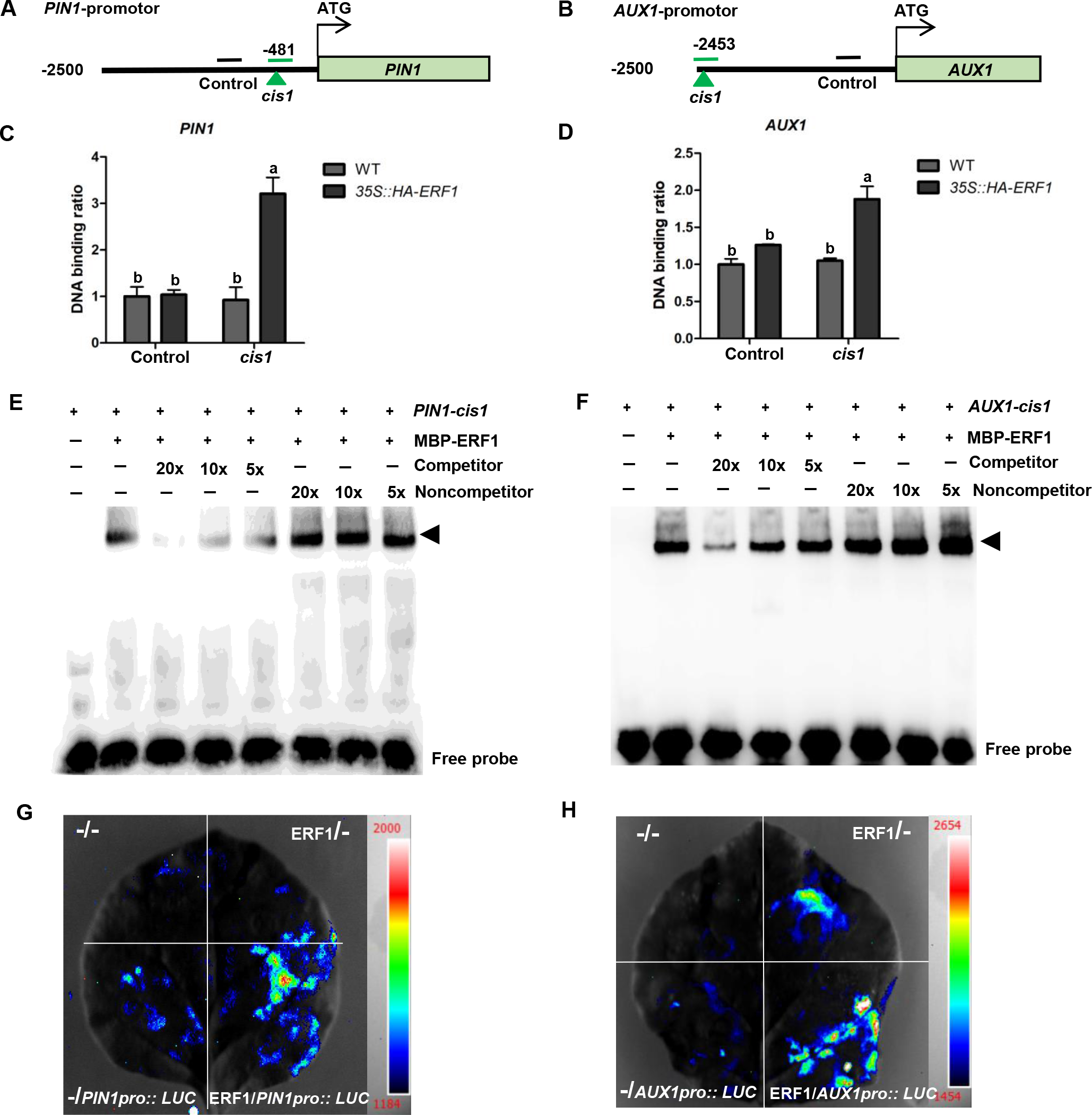
ERF1 activates *PIN1* and *AUX1* expression by directly binding to its promoter. (A and B) Schematic illustration of GCC-box in the *PIN1* and *AUX1* promoters. The number and green triangles represent the location of the GCC-box in the *PIN1* and *AUX1* promoters. The green lines represent the region with the GCC-box amplified in qPCR analysis, while the black lines mark the control region without the GCC-box. (C and D) ChIP qPCR assay. The binding of ERF1 to the *PIN1* and *AUX1* promoters *in planta* was assayed in 10-day-old wild type and *35S::HA-ERF1* transgenic plants with anti-HA antibodies. The *PIN1* or *AUX1* promoter region without a GCC-box was used as a control. Values are the mean ± SD (n=3 replicates). Different letters indicate significant differences by one- way ANOVA (P < 0.05). (E and F) EMSA. The biotin-labelled probes of a 26 bp *PIN1* or *AUX1* promoter fragment containing the GCC-box were incubated with MBP-ERF1 protein. Unlabelled probes were used as competitors, and mutated probes were used as noncompetitors. ERF1-dependent mobility shifts were competed by the competitor probe in a dose-dependent manner (5 x, 10 x, 20 x). Shifted bands are indicated by arrows. (G and H) Transient transactivation assays in tobacco leaves. The effector (pRI101/*ERF1*) and reporter (pGreenII 0800/*PIN1pro::LUC* and pGreenII 0800/*AUX1pro::LUC*) constructs were cotransfected into tobacco leaves, and the relative LUC fluorescence intensity was detected by the luciferase assay system as described in the Methods. “-/-” represents cotransfected pRI101 and pGreenII 0800 empty plasmids, “-/*PIN1pro::LUC*” and “-/*AUX1pro::LUC*” represent pRI101 and the reporter constructs, and “ERF1/-” represents the effector and pGreenII 0800 plasmids, which served as negative controls. “ERF1/*PIN1pro::LUC*” and “ERF1/*AUX1pro::LUC*” represent cotransfected effector and reporter constructs.

### ERF1 alters the expression of *PIN1* and *AUX1* in LRP

Given that ERF1 is involved in LR emergence by regulating the expression of *PIN1*, we investigated the transcript level of *PIN1* in *erf1-2*, WT, and OX-4 lines crossed with the *PIN1pro::GUS* reporter line. As shown in Figure 5A, the transcript level of *PIN1*, as indicated by the GUS signal, was significantly decreased in the *erf1-2* mutant but increased in the OX line compared with WT at stages I-VIII of LR formation, which is consistent with the qRTLPCR results (Figure S3A). Moreover, the intensity of GUS staining was significantly increased in the WT and OX-4 lines in response to IAA, while the *erf1-2* mutant displayed no apparent change (Figures 5A and 5B). Strong GUS signals were observed in the epidermis, cortex, and endodermis cells around the LRP of the OX-4 line (Figure 5A), similar to the *DR5::GUS* pattern in the OX line (Figure 2A).

**Figure 5.**
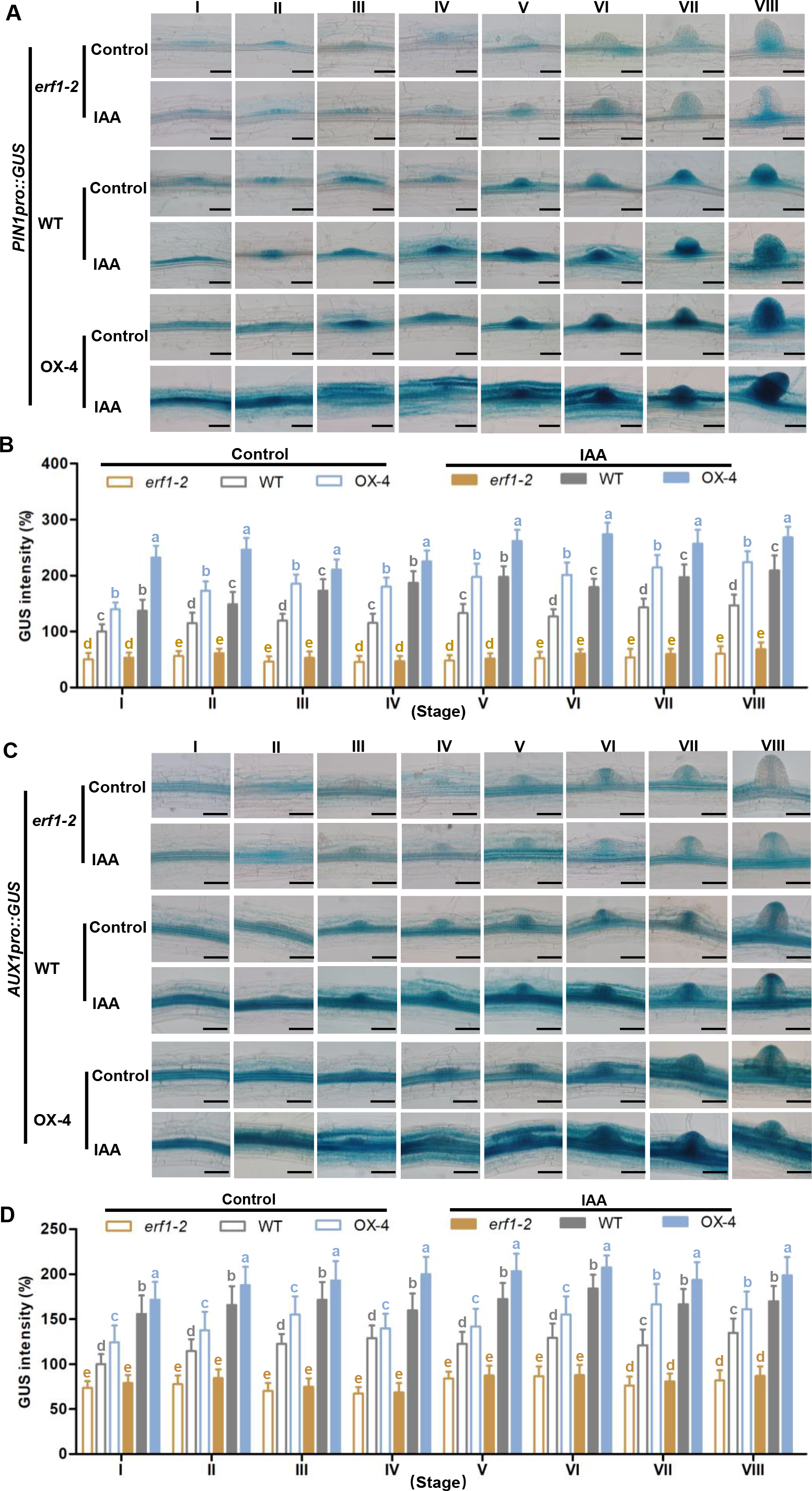
**ERF1 directly upregulates the expression of *PIN1* and *AUX1 in planta*** (A-D) Transcription of *PIN1* and *AUX1* in LRP. Seeds of *erf1-2*, WT, OX-4 lines in the *PIN1pro::GUS* or *AUX1pro::GUS* background were germinated on MS medium for 10 days, and then the seedlings were transferred to MS liquid without (Control) or with 200 nM IAA for 6 hours. Seedlings were incubated in GUS solution for 5 hours in (A) and 2 hours in (C) before the photographs of LRP from stage I to VIII were taken with a microscope. Bar=50 μm. The GUS signal intensity of *PIN1pro::GUS* (B) and *AUX1pro::GUS* (D) was quantified with ImageJ software. Values are the mean ± SD (20 images/genotype). Different letters indicate significant differences by one-way ANOVA (P < 0.05).

Additionally, the *PIN1pro::PIN1-GFP* reporter was introduced into the *erf1-2*, WT, and OX-4 lines, and the PIN1-GFP signals exhibited a similar pattern to that of the *PIN1Pro::GUS* reporter (Figure S4), indicating that ERF1 directly upregulates *PIN1* and enhances auxin transport during LR formation.

We also tracked the expression of *AUX1* in *erf1-2*, WT, and OX-4 lines by crossing with the *AUX1pro::GUS* reporter line. A lower GUS signal was present in the stele and LRP of the *erf1-2* mutant than in WT under normal conditions, whereas a higher GUS signal was observed in the OX-4 line (Figures 5C and 5D). Similarly, more intense GUS staining was observed in the primary root tips of OX-4 (Figure S5). Moreover, the transcript level of *AUX1* was upregulated in LRP and primary root tips of WT and OX-4 lines when subjected to IAA treatment, but no significant difference was observed in the *erf1-2* mutant (Figures 5C, 5D and S5). These results suggest that ERF1 increases *AUX1pro::GUS* expression in primary root tips, resulting in enhanced basipetal auxin transport.

### ERF1 directly represses the transcription of *ARF7*

Auxin signaling plays a vital role in LR formation. To investigate the possibility of ERF1 being involved in auxin signaling in LR emergence, we checked key genes related to auxin signaling modules. qRTLPCR results showed that *ARF6*, *ARF7*, *ARF19*, *LBD16*, and *LBD29* were upregulated in the *erf1-2* mutant and downregulated in the OX-4 line compared with WT plants. Moreover, IDA-HAE/HSL2 signaling module components were significantly repressed in the OX-4 line. Meanwhile, the transcript abundance of *XTR6*, *EXP17*, and *PG*, which are related to CWR, was enhanced in the *erf1-2* mutant and attenuated in the OX-4 line (Figure S6). These results suggest that ERF1 inhibits LR emergence, probably by modulating the expression of the key components related to auxin signaling and CWR.

We further showed that ERF1 was able to directly repress the expression of *ARF7* by binding to its promoter, as evidenced by ChIPLqPCR, electrophoretic mobility shift, and transient expression assays (Figures 6A-D).

**Figure 6.**
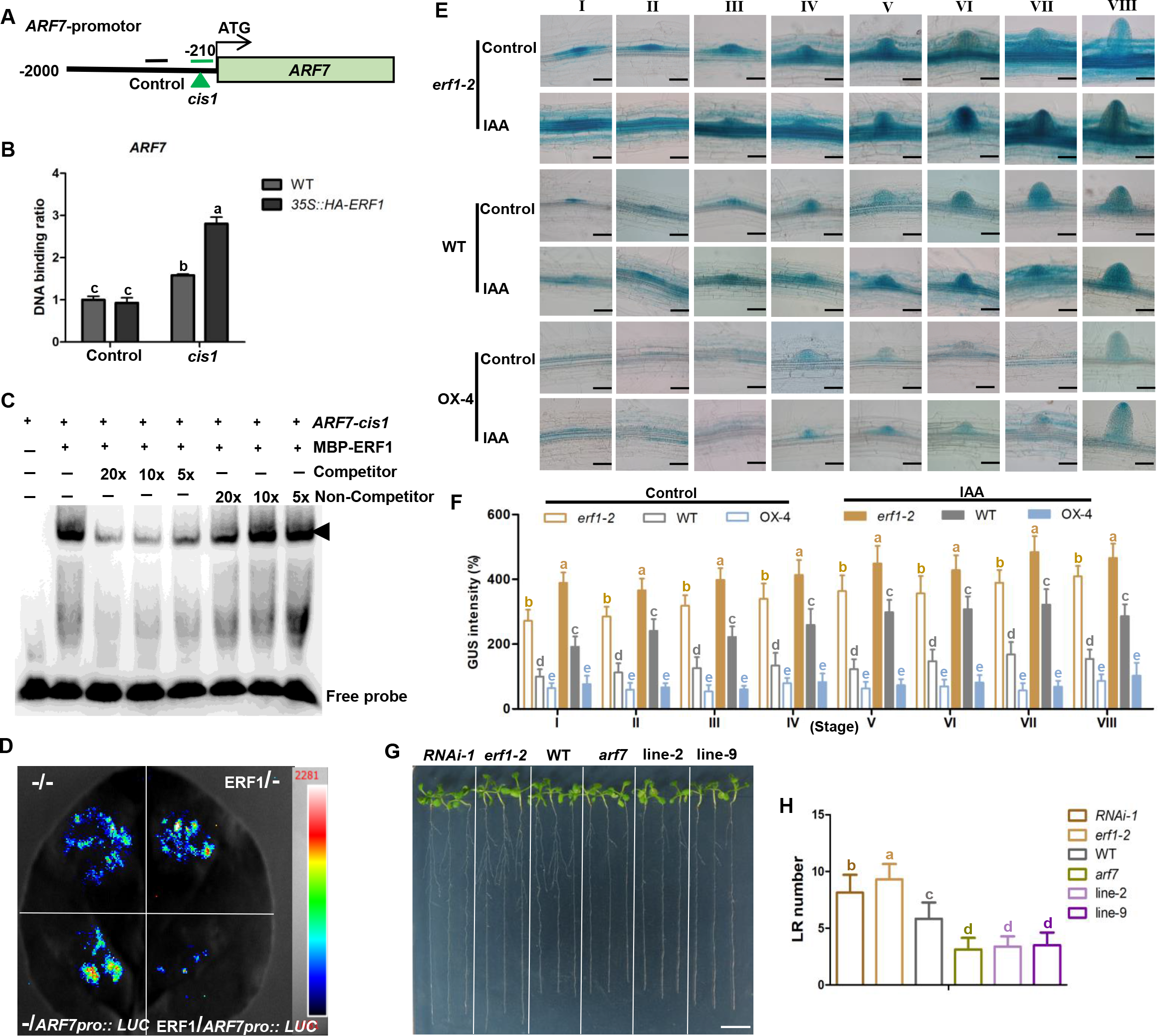
ERF1 represses the expression of *ARF7* by directly binding to its promoter. Schematic illustration of the GCC-box in the *ARF7* promoter. The number and green triangle represent the location of the GCC-box in the *ARF7* promoter. The green line represents the region with the GCC-box amplified in qPCR analysis, while the black line marks the control region without the GCC-box. (A) ChIP qPCR assay. The binding of ERF1 to the *ARF7* promoter *in planta* was assayed in 10-day-old wild-type and *35S::HA-ERF1* transgenic plants with anti-HA antibodies. An *ARF7* promoter region without a GCC-box was used as a control. Values are the mean ± SD (n=3 replicates). Different letters indicate significant differences by one-way ANOVA (P < 0.05). (B) EMSA. The biotin-labelled probe of a 26 bp *ARF7* promoter fragment containing the GCC-box was incubated with MBP-ERF1 protein. An unlabelled probe was used as a competitor, and a mutated probe was used as a noncompetitor. ERF1-dependent mobility shifts were competed by the competitor probe in a dose-dependent manner (5 x, 10 x, 20 x). Shifted bands are indicated by arrows. (C) Transient transactivation assays in tobacco leaves. The effector (pRI101/*ERF1*) and reporter (pGreenII 0800/*ARF7pro::LUC*) constructs were cotransfected into tobacco leaves, and the relative LUC fluorescence intensity was detected by the luciferase assay system as described in the Methods. “-/-” represents cotransfected pRI101 and pGreenII 0800 empty plasmids, “-/*ARF7pro::LUC*” represents pRI101 and the reporter construct, and “ERF1/-” represents the effector and pGreenII 0800 plasmids, which served as negative controls. “ERF1/*ARF7pro::LUC*” represents cotransfected effector and reporter constructs. (E and F) *ARF7* expression in LRP. Seeds of *erf1-2*, WT, OX-4 lines in the *ARF7pro::GUS* background were germinated on MS medium for 10 days, and then the seedlings were transferred to MS liquid with or without 200 nM IAA for 6 hours. Seedlings were incubated in GUS solution for 5 hours before the photographs of stages I to VIII were taken by microscope (E). Bar=50 μm. The GUS signal intensity (F) was quantified with ImageJ software. Values are the mean ± SD (20 images/genotype). Different letters indicate significant differences by one-way ANOVA (P < 0.05). (G and H) LR phenotype of WT, *RNAi-1*, *erf1-2*, *arf7*, line-2 (*RNAi-1 arf7*, #2), and line-9 (*erf1-2 arf7*, #9) lines. Seeds were germinated on MS medium for 7days, and then transferred to MS medium and grown vertically for 5 days before photographs were taken (G) and the LR number was counted (H). Values are mean ± SD (n=3 replicates, 30 seedlings/replicate). Different letters indicate significant differences by one-way ANOVA (P < 0.05). Bar=1 cm.

To determine whether the spatiotemporal expression pattern of *ARF7* was altered by ERF1, *ARF7pro::GUS* was introduced into *erf1-2*, WT, and OX-4 lines by crossing. Under normal conditions, a higher expression level of *ARF7pro::GUS* was observed in the LRP of the *erf1-2* mutant than in WT, but the OX-4 line displayed the opposite results. In response to auxin, the *ARF7pro::GUS* signal was induced in the *erf1-2* and WT lines, whereas no significant change was observed in the OX-4 line compared to the normal condition (Figures 6E and 6F). Together, these results show that ERF1 directly downregulates the expression of *ARF7*, consistent with the qRTLPCR results (Figure S6G).

To confirm the significance of the regulation of *ARF7* by ERF1, *RNAi-1 arf7* (line-2) and *erf1-2 arf7* (line-9) double mutants were generated by crossing. Fewer LR numbers were shown in the *arf7* line compared to WT, which is consistent with a previous study.^45^ However, the *RNAi-1 arf7* and *erf1-2 arf7* double mutant lines both exhibited similar LR emergence as *arf7*, suggesting that *ERF1* is epistatic to *ARF7* (Figures 6G and 6H).

### ERF1 modulates LR emergence in response to abiotic stress

*ERF1* was strongly responsive to salt, ethylene, and JA.^42, 46^ Here, our work further demonstrated that *ERF1* was also expressed in LRP at stages I-VII, endodermal, cortical, and epidermal cells adjacent to LRP as well as pericycle and stele cells and was strongly induced by mannitol, NaCl, ABA, IAA, and ACC treatments (Figures S7A and S7B). Intriguingly, the expression of *ERF1* decreased with LRP development, and it was barely detected in emerged LRs. These data suggest that ERF1 may play an important role in LR development in response to environmental cues.

To confirm the role of *ERF1* in LR emergence in response to stress, we transferred 7-day-old seedlings of *erf1*, WT, and OX lines grown on normal medium to MS medium with or without NaCl and mannitol. All the treatments produced similar results: the OX-2 and OX-4 lines had fewer LRs than the WT, whereas the *RNAi-1* and *erf1-2* mutants showed the opposite results (Figures S7C and S7D).

Similar results were observed in LR density (Figure S7E). However, no significant difference was displayed in *RNAi-1* and *erf1-2* mutants under NaCl or mannitol treatments compared with normal conditions, but salt and mannitol stress suppressed LR emergence of WT and OX lines compared to that of the control (normal condition). These results support the potential role of *ERF1* in regulating LR emergence in response to salt and mannitol stresses.

## Discussion

LRs are key constituents of plant root system architecture and sense environmental cues. In this study, we show that ERF1 functions as a negative regulator of LR emergence (Figure 1). *ERF1* is responsive to various environmental and hormonal signals (Figure S7). ERF1 enhances auxin transport by directly upregulating the expression of *PIN1* and *AUX1*, leading to auxin accumulation with altered distribution in the endodermal, cortical, and epidermal cells overlying LRP (Figures 2-5). Then, excessive auxin activates *ERF1* to form a positive feedback loop, and ERF1 directly represses *ARF7* transcription, thus affecting the expression of CWR genes and ultimately inhibiting LR emergence (Figure 6). Our results reveal that ERF1 integrates environmental signals into auxin signaling to adaptively regulate LR emergence in response to environmental fluctuations. A working model is proposed in Figure 7.

**Figure 7.**
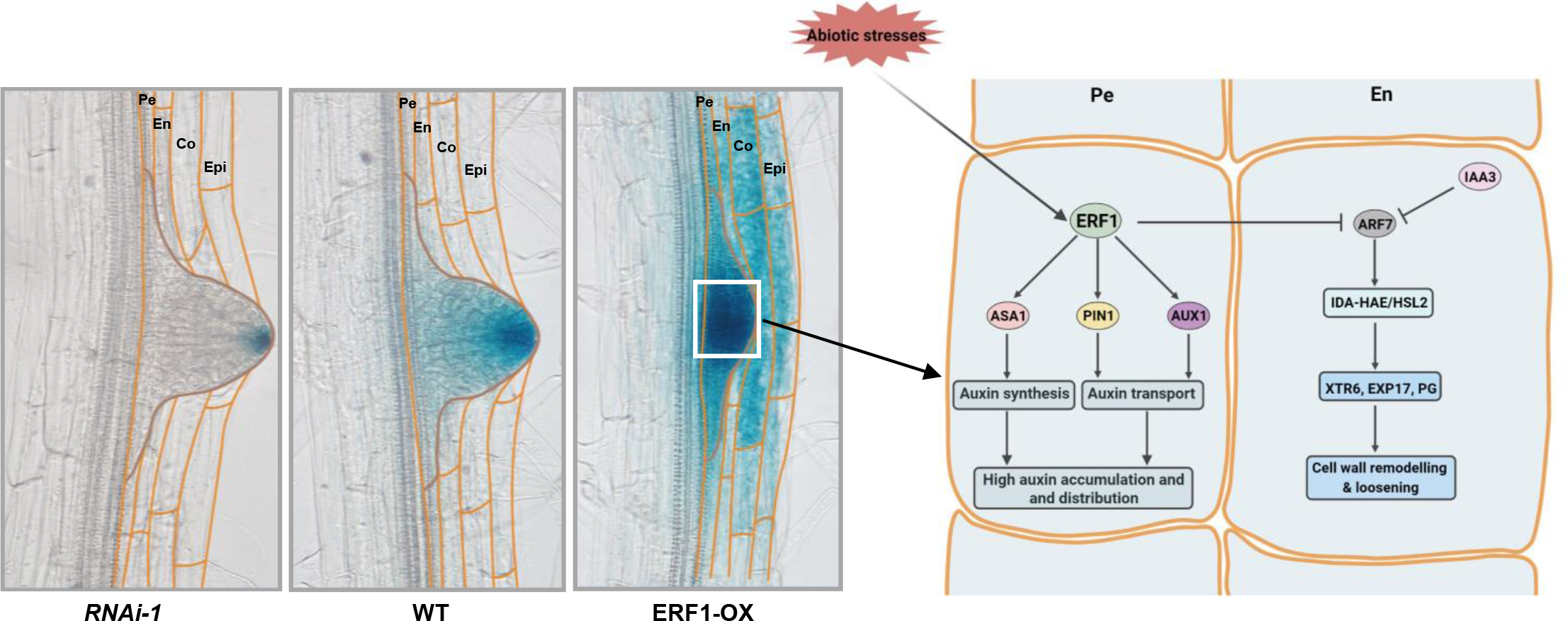
A working model for ERF1-mediated inhibition of LR emergence. ERF1 is responsive to various environmental and hormone signals and integrates these signals into auxin signaling. ERF1 enhances auxin biosynthesis by upregulating *ASA1* expression and increases auxin transport by upregulating *PIN1* and *AUX1* transcription, resulting in the perturbed distribution of high local auxin accumulation in the epidermis, cortex and endodermis, which leads to the inhibition of LR emergence. Additionally, ERF1 regulates LR emergence by repressing *ARF7* expression, which alters the expression of CWR genes through the IDA-HAE/HSL2 signaling module. Pe, pericycle; En, endodermis; Co, cortex; Epi, epidermis.

### ERF1 promotes auxin transport and alters the auxin distribution pattern

Auxin efflux carriers PIN-FORMEDs (PINs) and influx carriers AUX1/LAXs are involved in polar auxin transport across the plasma membrane to regulate LR development.^47, 48^ Previous studies showed that the amount of endogenous IAA and polar auxin transport (PAT) activity are reduced in the *pin1* mutant.^49, 50^ Meanwhile, the expression pattern of *PIN1* in the root meristem is directly related to the difference in endogenous auxin levels.^51^ The high-affinity auxin transport carrier AUX1 mediates auxin basipetal transport from the root tip to the elongation zone through the LRC and epidermal layers.^25, 52^ AUX1 and LAX3 are also involved in auxin signaling that activates *LBD16*/*ASL18* to control LR initiation and formation.^53^ Recently, SUE4, a PIN1-interacting membrane protein, was shown to interferes rootward auxin transport and reduce auxin levels in the root tip, resulting in faster elongation of the primary root.^54^ We demonstrated that downregulating *PIN1* and *AUX1* expression in *RNAi-1* and *erf1-2* mutants, contributed to decreased auxin transport and promoted

LR emergence (Figures 1B, 2C and 2D). Nevertheless, NPA treatment results in severe inhibition of LR formation by blocking LRP initiation.^32, 55^ NPA synergistically inhibits both PIN- and ABCB-based major auxin transport and probably interferes with critical phosphorylation events.^56^ NPA treatment inhibited LR formation in all three genotypes (Figures 3A-3C), although DR5-GUS signals were significantly reduced in WT and OX. However, the mechanism by which NPA inhibits auxin transport is inconsistent with that by downregulating *PIN1* in *erf1* mutants. Perhaps the modest decrease in auxin transport enhances the response to auxin in LRP of the *erf1* mutant, increasing *ARF7* expression, thus enhancing the expression of CWR genes and ultimately promoting LR emergence, which needs further investigation.

High auxin levels have been shown to inhibit cell elongation and promote LR formation. Interestingly, a previous study showed that high auxin-regulated inhibition of LR initiation in *Arabidopsis* is linked to reduced cell length.^57^ Meanwhile, recent evidence showed that high auxin concentrations (0.1 µM NAA) resulted in a reduced rate of LR initiation in maize, which was associated with inhibition of pericycle cell elongation.^58^ In our study, ERF1-mediated high auxin accumulation did not affect LR initiation but did affect emergence (Figure 1F). We assumed that the high auxin level in ERF1-OX lines probably did not enhance the frequency of auxin maxima required for LR initiation.

Auxin distribution is regulated by changing the expression, localization, or activity of auxin transporters. As previously shown, the process of LR formation involves dynamic changes in PIN-driven auxin distribution.^13, 49^ The differences in auxin levels in cell distribution caused by subcellular relocalization of PIN1 directly affect root growth.^59^ Environmental and endogenous signals can be integrated to regulate auxin distribution by affecting auxin transport.^30, 60^ Moreover, the ectopic overproduction of auxin in the stele, endodermis, and epidermis by expressing cell type-specific markers results in the inhibition of the primary roots.^33, 61^ The endodermis plays a key role in coordinating LR emergence.^5, 10, 62^ The endodermis serves as an important site for ABA-mediated inhibition of LR development in response to salt stress.^63^ Moreover, auxin activity in the endodermis coordinates the release of the restraints that inhibit meristematic activity of the pericycle.^64^ We reveal that ERF1 promotes local auxin accumulation in the endodermis, cortex, and epidermis surrounding LRP (Figure 2A), which might perturb the normal spatiotemporal auxin pattern for LR emergence. It is worth investigating whether the ERF1-mediated endodermal response inhibits LR emergence.

### ERF1 is involved in auxin signaling to regulate LR emergence

LR formation is a complex process that integrates multiple auxin signaling modules. Among those, the signaling module of SHY2/IAA3-ARF7 in the endodermis and SLR/IAA14-ARF7 in the cortex and epidermis both confer LRP organogenesis and outgrowth by regulating the separation of cells.^65^ The auxin response factor ARF7 not only plays crucial roles in auxin signaling but also functions in ethylene responses. *ERF1* transcript levels in *arf7* and *arf7 arf19* double mutants were similar to those in the wild type when treated with ethylene,^66^ suggesting that *ARF7* acts downstream of *ERF1*, which is consistent with our results (Figures 6G and 6H). Recently, SUMO-mediated posttranslational modification of ARF7 was shown to disrupt IAA3 recruitment; thus, repressing auxin-responsive gene expression was associated with LR initiation in response to water availability.^67^ Moreover, *PP2C. D1* expression is specifically activated by high levels of auxin via ARF7, leading to restrained cell elongation in the hypocotyl through PP2C. D1-mediated dephosphorylation of H^+^- ATPase.^68^ Whether the underlying mechanism by which ERF1-mediated high auxin accumulation alters cell wall acidification and suppresses cell expansion to affect LR emergence awaits further investigation.

The IDA-HAE/HSL2 signaling module, in which all components are expressed in the cells overlying the new LRP, is involved in regulating the expression of the CWR genes *XTR6*, *EXP17*, and *PG*.^41^ The expression levels of *IDA*, *HAE*, and *HSL2* were blocked in the *arf7* mutant and significantly reduced in the *arf19* mutant, suggesting that the activation of receptor genes (*HAE*, *HSL2*) is dependent on the upregulation of *ARF7* by an auxin-dependent cascade.^41^ Notably, the transcript abundance of *ARF7*, *IDA*, *HAE* and *XTR6*, *EXP17*, *PG* was repressed in the *ERF1* overexpression line, suggesting that ERF1 directly represses *ARF7* expression, thus affecting the expression of IDA-HAE/HSL2 signaling module components and CWR enzyme-encoding genes to inhibit LR emergence, which is in agreement with our results that ERF1 directly represses *ARF7* expression (Figures 6 and S6G). A number of studies have confirmed the significance of IDA-HAE/HSL2 signaling downstream of auxin-governed LR formation by regulating CWR genes. The mitogen-activated protein kinases MKK4/MKK5-MPK3/MPK6 were shown to function downstream of the IDA-HAE/HSL2 signaling module to regulate LR emergence by promoting the expression of CWR genes.^69^ The receptor-like protein kinase MUS, which is dependent on the functions of ARF7 and ARF19, controls LR formation in the early stages by regulating CWR.^70^

### ERF1 integrates stress signals into auxin signaling pathways to adaptively regulate LR emergence

Plants have evolved a set of mechanisms to cope with fluctuating environments. Among them, optimizing root architecture to adapt to adverse environments is a common strategy to increase survival. It is well known that auxin is a key mediator for plants to integrate environmental signals. In our study, we show that *ERF1* is responsive to various environmental and hormone signals and thus can integrate these signals into auxin transport and auxin signaling in addition to auxin biosynthesis to regulate LR emergence, which plays an important role in regulating the root system to adapt to changing environments. For example, when roots sense salt stress, plants may respond to minimize salt uptake by halting the increase in root surface area.

Apparently, an effective way is to pause LR emergence. Under such circumstances, ERF1-mediated high auxin accumulation and distribution may serve as one of the mechanisms to temporarily suspend LR emergence. When salt stress diminishes, *ERF1* expression decreases, thus alleviating the inhibition of LR emergence caused by the abnormal distribution of high auxin accumulation and downregulated CWR factors and finally resuming root growth.

*ERF1* may be exploited to improve crop root system architecture. An ideal root system architecture of crops usually exhibits deeper primary roots with more LRs for maximizing nutrient absorption and escaping drought. Based on our findings and a previous study,^42^ downregulation of *ERF1* expression in the roots would produce longer primary roots with more LRs in *Arabidopsis*. Whether this mechanism is conserved in crops awaits further exploration.

## Limitations of the study

Although our results have shown that ERF1 promotes local auxin accumulation by upregulating *PIN1* and *AUX1*, resulting in excessive auxin accumulation in LRP and the endodermis, cortex, and epidermis overlying LRP (Figure 2A), we acknowledge that more research needs to be performed to investigate the underlying mechanism by which the disturbed spatiotemporal auxin patterns inhibit LR emergence. *ERF1pro::GFP* will be used to analyze the localization of ERF1 in cell layers in future.

## Supporting information

ERF1-Auxin

## Acknowledgments

This study was supported by grants from the National Natural Science Foundation of China (31900230 to P.Z.), China Postdoctoral Science Foundation (2020T130634 and 2019M652200 to P.Z.) and Youth Innovation Foundation of University of Science and Technology of China (WK2070000186 to P.Z.).

## Author Contributions

P.Z., J.Z., and C.X. designed the experiments. P.Z., J.Z., Y.C., Z.Z., G.W., J.M. performed the experiments and data analyses. P.Z. wrote the manuscript. P.Z., S.T., and C.X. revised the manuscript. P.Z. and C.X supervised the project.

### Declaration of Interests

The authors declare no competing interests.

## Supplemental Information

Figure S1. Verification of *RNAi-1*, *erf1-2* and *ERF1* overexpression lines (OX-2, OX- 4). Related to Figure 1.

Figure S2. Quantification of GUS signal intensity in response to NPA. Related to Figure 3D.

Figure S3. ERF1 alters the expression of the genes involved in auxin transport. Related to Figure 2.

Figure S4. Protein levels of PIN1-GFP in LRP and primary root tips. Related to Figure 4 and Figure 5.

Figure S5. ERF1 upregulates the expression of *AUX1pro::GUS* in primary root tips. Related to Figure 4 and Figure 5.

Figure S6. ERF1 alters the expression of the genes associated with auxin signaling and CWR. Related to Figure 6 and Figure 7.

Figure S7. ERF1 integrates abiotic stress signals to regulate LR emergence. Related to Figure 1.

Table S1. Primers used in this study. Related to Figures 4, 5, 6, S3, and S6.

## STAR Methods

### Resource Availability

#### Lead Contact

Further information and requests for resources should be directed to and will be fulfilled by the Lead Contact, Chengbin Xiang.

#### Materials Availability

The materials generated in this study are available from the corresponding author.

#### Data and code availability

- All data reported in this paper will be shared by the lead contact upon request.
- This paper does not report original and new code.
- Any additional information required to reanalyze the data reported in this paper is available from the lead contact upon request.

### Experimental model and subject details

#### Plant materials and growth conditions

*Arabidopsis thaliana* ecotype Col-0 was used as the wild type, and all *Arabidopsis thaliana* mutants and transgenic plants used in this study were in the Columbia (Col-0) background. The *35Spro::ERF1*-RNAi (*RNAi-1*), *ERF1*-overexpressing plants (OX-2, OX-4), *ERF1pro::GUS*, and *35S::HA-ERF1* plants used in this study were previously described.^42^ The *ERF1* knockout mutant *erf1-2* was generated by the CRISPRLCas9 technique as described.^71^ Double mutants were generated by crossing plants with distinct genetic backgrounds.

For seedling growth, surface-sterilized *Arabidopsis* seeds were treated with 10% bleach for 15 minutes, cold-treated for 2-3 days at 4°C, and then grown on Murashige and Skoog (MS) medium with 1% sucrose for the indicated days at 22 °C and 70-80% relative humidity under a 16/8-h light/dark photoperiod.

### Method Details

#### Analysis of the LR number and LRP number

Seeds were germinated on MS medium for 5 days, and then seedlings were transferred to MS medium for 8 days. The LR number was counted every other day for 8 days, and the LR number cm^-1^ was measured at the indicated time points. LRP was counted by using a microscope with a camera (HiROX).

#### GUS staining

GUS staining was performed as described.^72^ Transgenic plants (*ERF1pro::GUS* or *DR5::GUS*) with or without stress treatment were immersed in staining solution at 37°C in the dark, destained with 30%, 70%, and 100% ethanol and rehydrated with 100%, 70%, and 30% ethanol to remove chlorophyll. Samples were photographed using a microscope with a camera (HiROX).

#### Measurement of auxin polar transport in *Arabidopsis* roots

Rootward and shootward auxin transport was performed as previously described.^73^ Seven-day-old seedlings were grown on MS medium under long-day conditions (16/8-h light/dark photoperiod). ^3^H-IAA agar blocks were placed at the rootLshootjunction of seedlings for rootward auxin transport and at the root tip for the shootward auxin transport assay and then incubated for 5 hours in darkness. Approximately 10 mm of the apical root tip was harvested and quantified by using a scintillation counter.

To calculate the amount of ^3^H-IAA transported: fmol = ^3^H-d.p.m. × 0.0173.

#### RNA isolation and qRTDPCR analyses

Total RNA samples were isolated from seedling roots with RNA-easy isolation reagent (Vazyme, #r701-02). Reverse transcription-PCR was performed using a HiScript III 1st strand cDNA synthesis kit (Vazyme, #r323-01). Real-time qRTLPCR was carried out using ChamQ SYBR qPCR Master Mix (Vazyme, #q311-02) and the specific primers listed in Table S1. The transcript levels of different genes were normalized to that of *UBIQUITIN5* (*UBQ5*) in qRTLPCR analysis. Three independent biological replicates were performed for each experiment, and the values presented represent the means ± SDs.

#### Chromatin immunoprecipitation (ChIP) assay

The ChIP assay was carried out as described previously.^74^ Ten-day-old *35S::HA-ERF1* transgenic seedlings and wild type plants were used for the ChIP assay. The immunoprecipitated DNA fragments were analyzed by qPCR with specific primers, and the enrichment in *35S::HA-ERF1* transgenic seedlings and wild type plants was calculated. β*-Tubulin8* and an approximately 200 bp promoter fragment without a binding site were used as negative controls.

#### Electrophoretic mobility shift assay

An electrophoretic mobility shift assay (EMSA) was conducted using a LightShift™ EMSA Optimization and Control Kit (20148×) (Thermo Fisher Scientific, Waltham, USA) as described previously.^75^ The coding region of *ERF1* was cloned into the pMAL c2x vector, and the MBP-ERF1 fusion protein was expressed in the *E. coli* Rosseta2 strain. Twenty-six bp free probes containing the GCC-box, competitor and noncompetitor probes (mutated) were obtained by slowly cooling from 98°C to 25°C for 10 min. EMSA reaction mixtures were loaded on 6% polyacrylamide gel in 0.5 × TBE buffer electrophoresed at 4°C.

#### Transient expression assay in *Nicotiana benthamiana* leaves

A transient expression assay in tobacco leaves was conducted as described previously.^76^ The coding sequence of *ERF1* was cloned into the pRI101 vector. Approximately 2500 bp promoters of *PIN1*, *AUX1*, and *ARF7* were introduced into the pGreenII 0800-LUC vector. All constructs were electroporated into the *Agrobacterium* GV3101 strain by using MicroPulser Electroporation Systems (Bio-Rad, USA, CA) and then coinjected into *Nicotiana benthamiana* leaves with the infiltration solution (10 mM MES, 10 mM MgCl2, 150 mM acetosyringone, pH 5.6). Two days after injection, tobacco leaves were sprayed using Luc substrates (Xeno light^TM^ D-luciferin potassium salt, 1 mM).

### Quantification and statistical analysis

All data presented in this study are expressed as the mean values ± SD of at least three replicates. Statistical analyses were performed by SPSS v16.0 software (Chicago, IL, USA) based on one-way ANOVA, and significant differences were determined at P≤0.05.

## Notes

### Competing Interest Statement

The authors have declared no competing interest.

