## Supplementary material for "ERF1 inhibits lateral root emergence by promoting local auxin accumulation with altered distribution and repressing *ARF7* expression": ERF1-Auxin

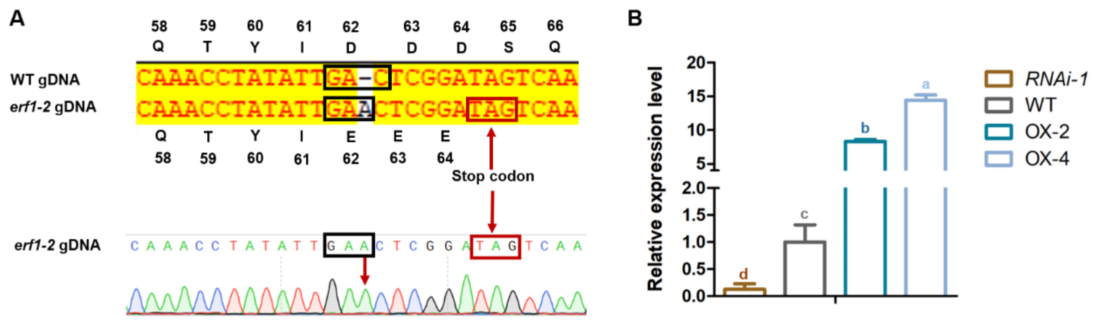


**Figure S1. Verification of *RNAi-1*, *erf1-2* and *ERF1* overexpression lines (OX-2, OX-4). Related to Figure 1.**

(A) Identification of CRISPR/Cas9-edited *ERF1* knockout mutant *erf1-2*. Sequencing results of *erf1-2* mutant showed a nucleotide insertion in the codon of the 62$^{th}$ amino acid resulting in the change the 65$^{th}$ codon from AGT to TAG.

(B) The expression level of *ERF1* in 7-day-old seedlings. Seeds of *RNAi-1*, WT, OX-2 and OX-4 lines were grown on MS medium for 7 days, then RNA was isolated from seedlings. The transcript level of *ERF1* was analyzed by qRT-PCR. Values are mean ± SD (n=3 replicates). Different letters indicate significant differences by one-way ANOVA ($P < 0.05$).

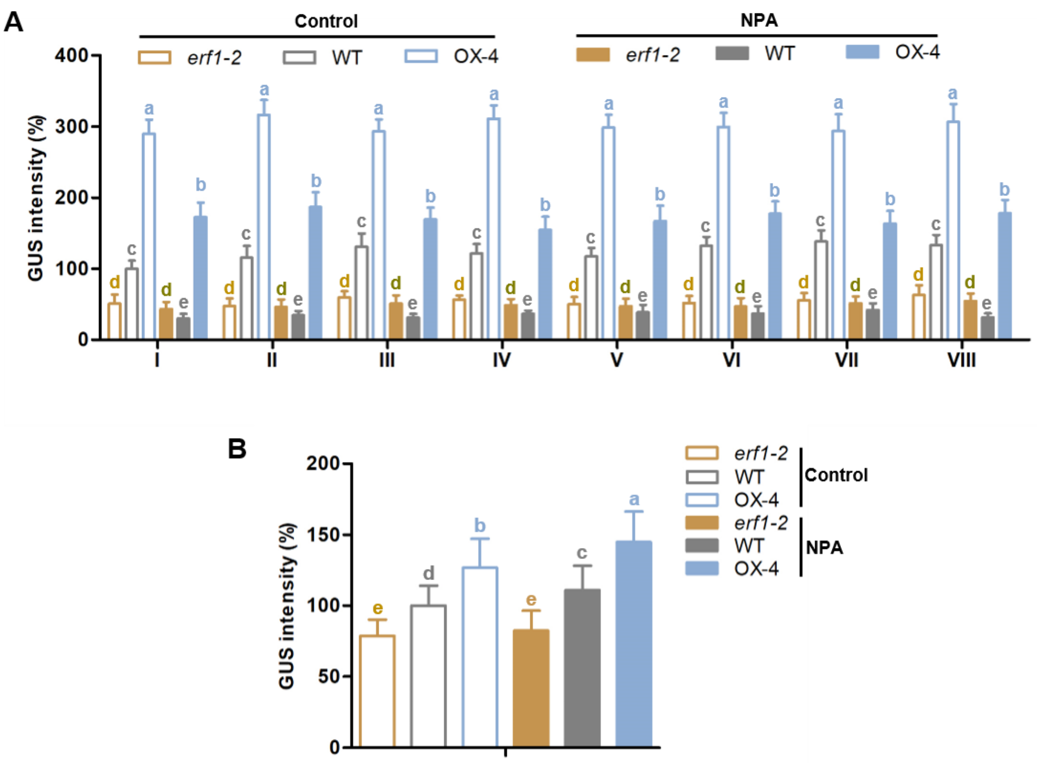


**Figure S2. Quantification of GUS signal intensity in response to NPA. Related to Figure 3D.**

(A and B) The relative GUS signal intensity of *erf1-2*, WT, and OX-4 lines in LRP (A) and root tips (B), as shown in (Figure 3D), was quantified with ImageJ software. Values are the mean ± SD (20 images/genotype). Different letters indicate significant differences by one-way ANOVA ($P < 0.05$).

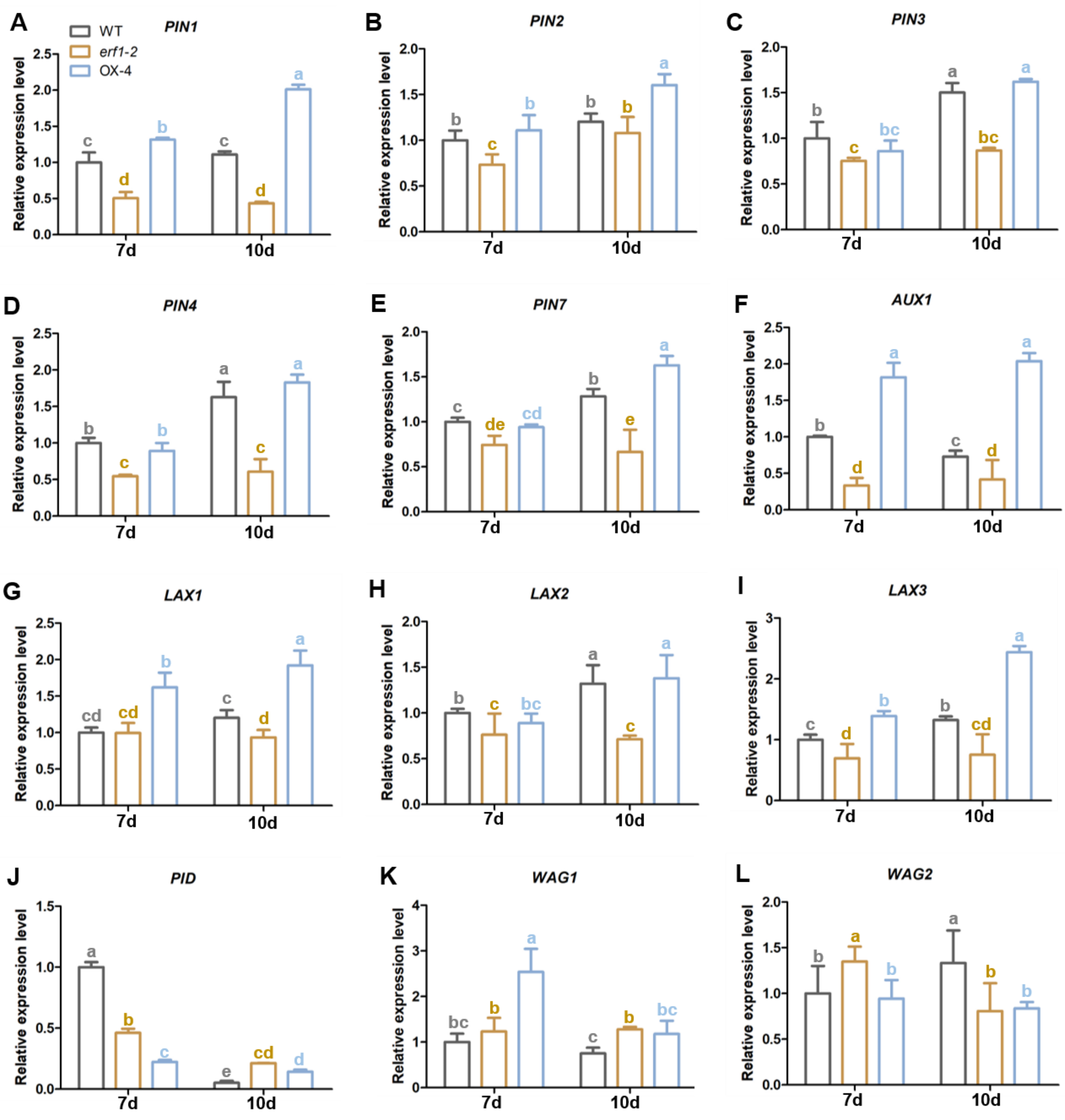


**Figure S3. ERF1 alters the expression of the genes involved in auxin transport. Related to Figure 2.**

(A-L) Seeds of *erf1-2*, WT, OX-4 lines were germinated on MS medium for 7, 10 days respectively, then RNA was isolated from roots. The transcript levels of the genes related to auxin transport (A-I) and AGCVIII kinase (J-L) were analyzed by qRT-PCR. Values are mean ± SD (n=3 replicates). Different letters indicate significant differences by one-way ANOVA (P < 0.05).

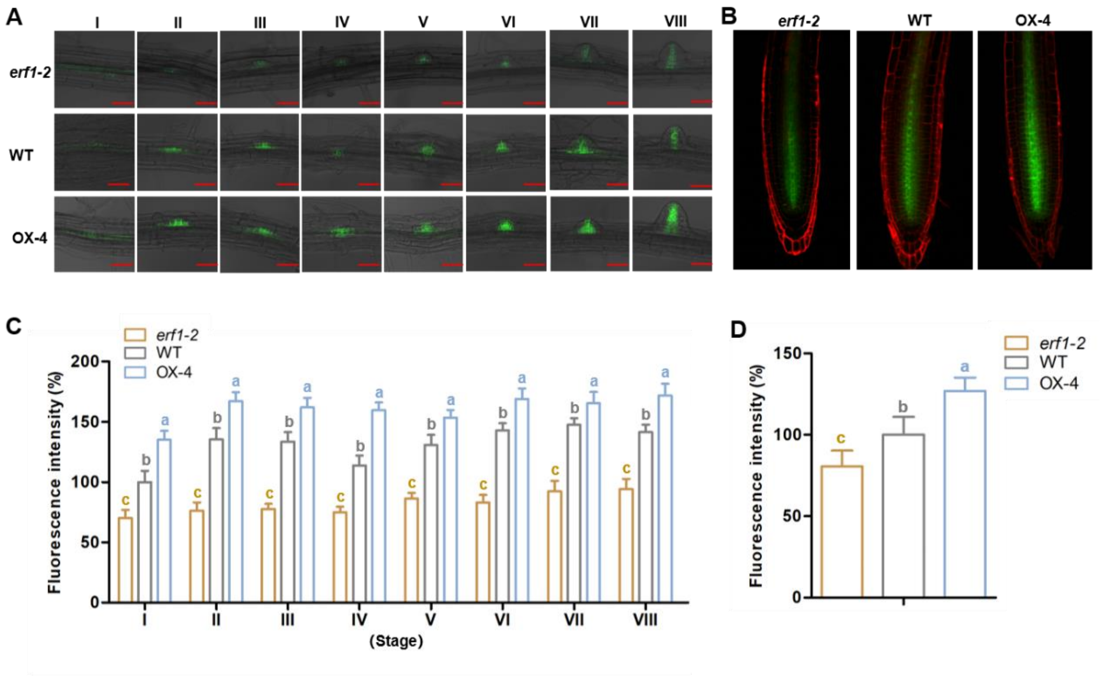


**Figure S4. Protein levels of PIN1-GFP in LRP and primary root tips. Related to Figure 4 and Figure 5.**

(A-D) Ten-day-old seedling roots of *erf1-2*, WT, and OX-4 lines in *PIN1pro::PIN1-GFP* background were visualized using a confocal laser scanning microscope. The photographs of LRP from stage I to VIII (A) and primary root tips (B) were taken. All images were acquired with identical confocal settings (AF488: lasers 2.6%, Pinhole 2.07 AU, Master Gain 528V, Digital Offset -24, Digital Gain 1.0; AF543: lasers 10%, Pinhole 2.07 AU, Master Gain 500V, Digital Offset -32, Digital Gain 1.0). Bar=50 µm. The GFP fluorescence intensity of LRP (C) and primary root tips (D) was quantified with ImageJ software. Values are the mean ± SD (20 images/genotype). Different letters indicate significant differences by one-way ANOVA ($P < 0.05$).

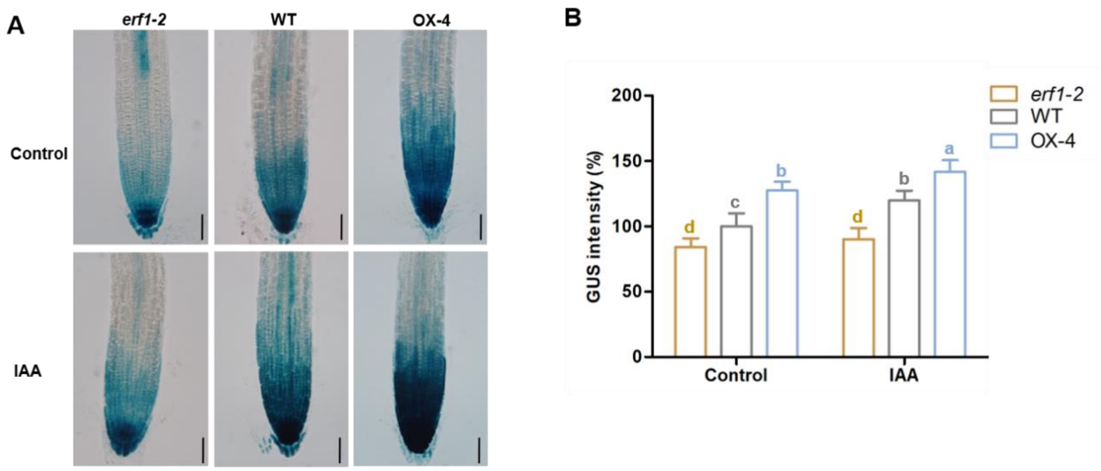


**Figure S5. ERF1 upregulates the expression of *AUX1pro::GUS* in primary root tips. Related to Figure 4 and Figure 5.**

(A and B) Seeds of *erf1-2*, WT, and OX-4 lines in *AUX1pro::GUS* background were germinated on MS medium for 10 days, then the seedlings were transferred to MS liquid without (Control) or with 200 nM IAA for 6 hours before seedlings were incubated in GUS solution for 2 hours before photographs of primary root tips were taken (A). Bar=50 μm. The GUS signal intensity (B) was quantified with ImageJ software. Values are the mean ± SD (20 images/genotype). Different letters indicate significant differences by one-way ANOVA ($P < 0.05$).

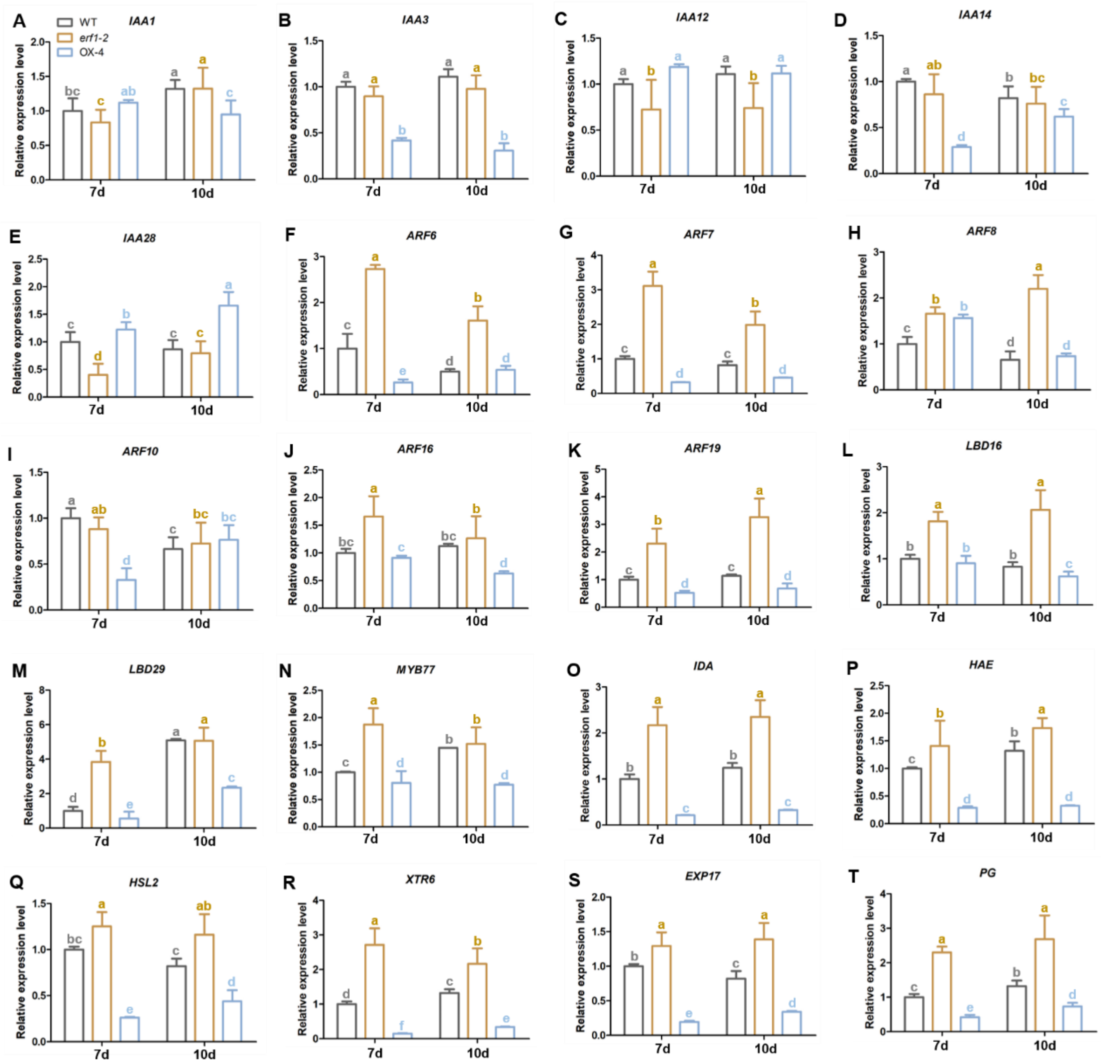

**Figure S6. ERF1 alters the expression of the genes associated with auxin signaling and CWR. Related to Figure 6 and Figure 7.**

(A-T) Seeds of *erf1-2*, WT, and OX-4 lines were grown on MS medium for 7, 10 days respectively, then RNA was isolated from roots. The transcript levels of genes-associated auxin signaling (A-N), IDA-HAE/HSL2 signaling (O-Q) and CWR (R-T) were analyzed by qRT-PCR. Values are mean ± SD (n=3 replicates). Different letters indicate significant differences by one-way ANOVA ($P < 0.05$).

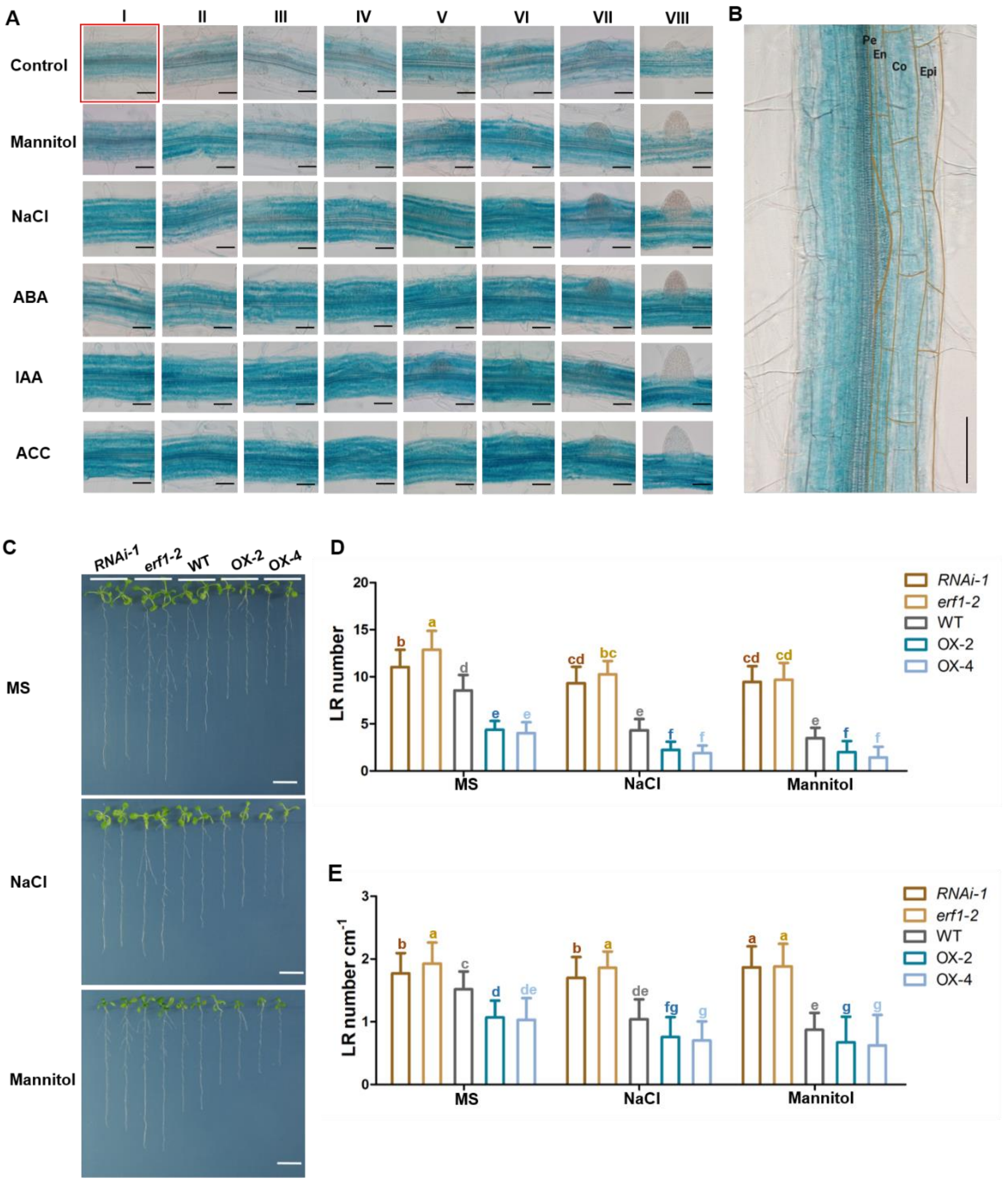


**Figure S7. ERF1 integrates abiotic stress signals to regulate LR emergence. Related to Figure 1.**

(A-B) Seven-day-old *ERF1pro::GUS* seedlings were transferred to MS liquid (Control) or MS liquid supplemented with 250 mM mannitol, 120 mM NaCl, 5 μM ABA, 200 nM IAA, or 1 μM ACC, respectively, and incubated for 6 hours. Afterwards, seedlings were incubated in GUS staining solution for 6 hours before the photographs of stages I to VIII were taken with a microscope (A). The enlarged picture in red box was shown in (B). Bar=50 μm.

(C-E) ERF1 regulates LR emergence under stress. Seeds of *RNAi-1*, *erf1-2*, WT, OX-2 and OX-4 lines were germinated on MS medium for 7 days respectively, then seedlings were transferred to MS medium with or without 75 mM NaCl, 200 mM mannitol for 5 days. Photographs were taken (C). LR number (D) was counted and LR number $cm^{-1}$ (E) were measured at day 5. Values are mean ± SD (n=3 replicates, 30 seedlings/replicate). Different letters indicate significant difference by one-way ANOVA ($P < 0.05$). Bar=1 cm.

**Table S1. Primers used in this study. Related to Figures 4, 5, 6, S2, S5 and S8.**

| Purpose | Primer | Sequence |
|---|---|---|
| For CHIP-PCR | *ARF7* cis1 P1 | TCGTATGCTTTGTTTGTCTCTCC |
| | *ARF7* cis1 P2 | CGTTAATCAGAAATGGGGAACTC |
| | *ARF7* control P1 | CTTCGATTGTCCTTGGTCATCC |
| | *ARF7* control P2 | GGAGAGACAAACAAAGCATACGA |
| | *PIN1* cis1 P1 | AGGGGATAAATTGATCAACCGAA |
| | *PIN1* cis1 P2 | TGGTTGATGGGTGCGTGGTGG |
| | PIN1 control P1 | CTCACACACACCCATCTCACAT |
| | *PIN1* control P2 | GATGATATAAGGCTCGATTTGATG |
| | *AUX1* cis1 P1 | AGAGGTGGGTTGGAGTCTTG |
| | *AUX1* cis1 P2 | ATACATAGAGCCTATGTATCTA |
| | *AUX1* control P1 | CTTCTCCTTCCCTTCTCTCTC |
| | *AUX1* control P2 | AGCTGTCACTTGTTTCTTCTC |
| For yeast-one hybrid | pAD/*ERF1* P1 | CGGCTAGCATGGATCCATTTTTAATTCAGT |
| | pAD/*ERF1* P2 | GCGTCGACTCACCAAGTCCCACTATTTTCAG |
| | pHIS2/*ARF7* cis1 LP | CGAGAAATTGGCGGCTCTGTCGGA |
| | pHIS2/*ARF7* cis1 RP | CGCGTCCGACAGAGCCGCCAATTTCTCGAGCT |
| | pHIS2/*PIN1* cis1 LP | CCGCACAAGGCCGCCTCTTTCACA |
| | pHIS2/*PIN1* cis1 RP | CGCGTGTGAAAGAGGCGGCCTTGTGCGGAGCT |
| | pHIS2/*AUX1* cis1 LP | CGCAACATTGCCGCCTTTACATAA |
| | pHIS2/*AUX1* cis1 RP | CGCGTTATGTAAAGGCGGCAATGTTGCGAGCT |

| | | |
|---|---|---|
| | *ARF7* cis1 +P1 | 5'biotinCCGAGAAATTGGCGGCTCTGTCGGAG-3' |
| | *ARF7* cis1 P2 | CTCCGACAGAGCCGCCAATTTCTCGG |
| | *ARF7* cis1 -P1 | CCGAGAAATTGGCGGCTCTGTCGGAG |
| | *ARF7* cis1 +P1-M | 5'biotinCCGAGAAATTGTCTACTCTGTCGGAG-3' |
| | *ARF7* cis1 P2-M | CTCCGACAGAGTAGACAATTTCTCGG |
| | *PIN1* cis1 +P1 | 5'biotinAGCGCACAAGGCCGCCTCTTTCACTA-3' |
| | *PIN1* cis1 P2 | TAGTGAAAGAGGCGGCCTTGTGCGCT |
| | *PIN1* cis1 -P1 | AGCGCACAAGGCCGCCTCTTTCACTA |
| | *PIN1* cis1 +P1-M | 5'biotinAGCGCACAAGGTAGACTCTTTCACTA-3' |
| | *PIN1* cis1 P2-M | TAGTGAAAGACTACTCCTTGTGCGCT |
| For EMSA assay | *AUX1* cis1 +P1 | 5'biotin-CTGCAACATTGCCGCCTTTACATAAA 3' |
| | *AUX1* cis1 P2 | TTTATGTAAAGGCGGCAATGTTGCAG |
| | *AUX1* cis1 -P1 | CTGCAACATTGCCGCCTTTACATAAA |
| | *AUX1* cis1 +P1-M | TTTATGTAAAGTAGACAATGTTGCAG |
| | *AUX1* cis1 P2-M | CTGCAACATTGTCTACTTTACATAAA |
| | pGreenII 62-SK/*ERF1* P1 | CGAGCTCGATGGATCCATTTTTAATTCAGTCC |
| | pGreenII 62-SK/*ERF1* P2 | AACTGCAGTCACCAAGTCCCACTATTTTCA |
| | pGreen0800/*ARF7* P1 | GCGTCGACTTGTTCTTTGTGATCGCATATGC |
| For transient assay | pGreen0800/*ARF7* P2 | AACTGCAGAATCTGAATCTGAGCTTATACAAA |
| | pGreen0800/*ARF7* P1 cis1 | TCGACCCGAGAAATTGGCGGCTCTGTCGGAGCTGCA |
| | pGreen0800/*ARF7* P2 cis1 | GCTCCGACAGAGCCGCCAATTTCTCGGG |
| | pGreen0800/*PIN1* P1 | CCCTCGAGCGCAACTACAACTGTAAATGAT |

| | | |
|---|---|---|
| | pGreen0800/*PIN1* P2 | GCGTCGAC GTTCGCCGGAGAAGAGAGAG |
| | pGreen0800/*PIN1* P1 cis1 | TCGAGAGCGCACAAGGCCGCCTCTTTCACT AG |
| | pGreen0800/*PIN1* P2 cis1 | TCGACTAGTGAAAGAGGCGGCCTTGTGCG CT C |
| | pGreen0800/*AUX1* P1 | CCCTCGAGGTGGGTTGGAGTCTTGAAGAC |
| | pGreen0800/*AUX1* P2 | CCAAGCTTGTTCTTCGTTATCTTTCCCGGT |
| | pGreen0800/*AUX1* P1 cis1 | GCAACATTGCCGCCTTTACATA |
| | pGreen0800/*AUX1* P2 cis1 | TATGTAAAGGCGGCAATGTTGC |
| | pRI101/*ERF1* P1 | GCGTCGACATGGATCCATTTTTAATTCAGT CC |
| | pRI101/*ERF1* P2 | CGAGCTCGTCACCAAGTCCCACTATTTTCA |
| For *ARF7pro::GUS* construction | pCAM1391Z/*ARF7* P1 | TGACCATGATTACGCCAAGCTTATCTCCAA ACTTCTGTCTTCAC |
| | pCAM1391Z/*ARF7* P2 | AAAACGACGGCCAGTGAATTCTTAAAAGA TACCATACATTCAGTG |
| For *PIN1pro::GUS* construction | pCAM1391Z/*PIN1* P1 | AACTGCAGTGTATCCACTTATCATTCCCAT T |
| | pCAM1391Z/*PIN1* P2 | GGAATTCTAGAACGACGAACAGTAACATG |
| For *AUX1pro::GUS* construction | pCAM1391Z/*AUX1* P1 | CCAAGCTTGTGGGTTGGAGTCTTGAAGAC |
| | pCAM1391Z/*AUX1* P2 | CGGGATCCGTTCTTCGTTATCTTTCCCGGT |
| | *UBQ5* P1 | AGAAGATCAAGCACAAGCAT |
| | *UBQ5* P2 | CAGATCAAGCTTCAACTCCT |
| | *ERF1* P1 | GCGGAGAGAGTTCAAGAGTCG |
| For qRT-PCR | *ERF1* P2 | GCTCCTCAAGGTACTGTTCTC |
| | *IAA12* P1 | GATGGAGTTGGTATAGGCAGAA |
| | *IAA12* P2 | AAAGAACATTTCCTCAAGCGTC |
| | *IAA28* P1 | AAAGCTCGACCAAAGAAACATC |

| | |
|---|---|
| *IAA28* P2 | GTGAAAGCTGTTGGTAGTTGTT |
| *IAA14* P1 | GCTCCTTTACCATGGGGAGT |
| *IAA14* P2 | TTGAACTTCTCCATTGCTCTTGG |
| *ARF10* P1 | GAGTTTCCATTCCACGGTACTA |
| *ARF10* P2 | ATCAGACAACAAAGACGGAGAT |
| *MYB77* P1 | CTTCTTAACGGTCGTACGGATA |
| *MYB77* P2 | GTTAGGACTCTCAGGACTCATG |
| *ARF19* P1 | GAACACGGTTAACCATATCAGC |
| *ARF19* P2 | GCTGTGACATCTGTAGTTGTTG |
| *LBD16* P1 | ATGTAAACCCTAACAATCCGGT |
| LBD16 P2 | ATTCCATCGTGACCGTATACTC |
| *LBD29* P1 | ACAGAGAGTAGTTACCACAACG |
| *LBD29* P2 | CCTGATTGAAAGTGTTCAGGTG |
| *IDA* P1 | ATGGCTCCGTGTCGTACGATG |
| *IDA* P2 | GAGGAAGAGAGTTAACAAAAGAG |
| *HAE* P1 | GCCTCTCTCGGTAACGTCAC |
| *HAE* P2 | GGATTCTGGTAACTCGCCTGA |
| *HSL2* P1 | CTCTGTTCCGCCTATTGTTCA |
| *HSL2* P2 | AATGTAGCCGTAGGATCCAGC |
| *XTR6* P1 | CCTGGATCTACATGGGACGA |
| *XTR6* P2 | CAGAGGATAGAGTAAGTGTGG |
| *EXP17* P1 | TCCACCGAACTTTGCTCAGGC |

| | |
|---|---|
| *EXP17* P2 | CTCACCCCCTCCTGCTACG |
| *AIR3* P1 | CCCTGAAGTTGTGTCTGTCTTC |
| *AIR3* P2 | CCTTGCTCCTATCAGTTTCCTA |
| *PGLR* P1 | CGGTGTTACGGTCAGTGGAATA |
| *PGLR* P2 | CTTTCCCAGTCCTCCAATGCT |
